# Role of graphene oxide in inhibiting the interactions between nucleoside diphosphate kinases -B and -C

**DOI:** 10.1101/2021.11.12.465807

**Authors:** Andrey O. Zaznaev, Isaac G. Macwan

## Abstract

During a heart failure, higher amount of nucleoside diphosphate kinase (NDPK) enzyme in the sarcolemma membrane inhibits the synthesis of second messenger cyclic adenosine monophosphate (cAMP), which is required for the regulation of the calcium ion balance for normal functioning of the heart. In a dependent pathway, NDPK normally phosphorylates the stimulatory guanosine diphosphate, GDP(s), to a guanosine triphosphate, GTP(s), on the heterotrimeric (α, β and γ subunits) guanine nucleotide binding protein (G protein), resulting in the stimulation of the cAMP formation. In case of a heart failure, an increased quantity of NDPK also reacts with the inhibitory GDP(i), which is converted to a GTP(i), resulting in the inhibition of the cAMP formation. Typically, the βγ dimer of the G protein binds with hexameric NDPK-B/C complex and receives the phosphate at the residue His266 from residue His118 of NDPK-B. It is known that NDPK-C is required for NDPK-B to phosphorylate the G protein. In this work, the interactions between NDPK-B and NDPK-C are quantified in the presence and absence of graphene oxide (GO) as well as those between NDPK-B and GO through stability analysis involving hydrogen bonds, center of mass (COM), root mean square deviation (RMSD), and salt bridges, and energetics analysis involving van der Waals (VDW) and electrostatic energies. Furthermore, the role of water molecules at the interface of NDPK-B and NDPK-C as well as between NDPK-B and GO is investigated to understand the nature of interactions. It is found that the adsorption of NDPK-B on GO triggers a potential conformational change in the structure of NDPK-B, resulting in a diminished interaction with NDPK-C. This is confirmed through a reduced center of mass (COM) distance between NDPK-B and GO (from 40 A□ to 30 A□) and an increased COM distance between NDPK-B and NDPK-C (from 50 A□ to 60 A□). Furthermore, this is also supported by fewer salt bridges between NDPK-B and NDPK-C, and an increased number of hydrogen bonds formed by the interfacial water molecules. As NDPK-C is crucial to be complexed with NDPK-B for successful interaction of NDPK-B with the G protein, this finding shows that GO can suppress the interactions between NDPK-B/C and G proteins, thereby providing an additional insight into the role of GO in the heart failure mechanism.

**AUTHOR SUMMARY:** We report a novel computational understanding of the interactions between the enzymes NDPK-B and NDPK-C with GO as a potential inhibitor to such interactions and its implications. These types of interactions can play influential roles in many biochemical processes including those that take place during heart failure. A second messenger called cAMP is needed for proper cardiac contraction through the actions of NDPK-B/NDPK-C. It is needed to study the interactions between NDPK-B and NDPK-C to control the synthesis of cAMP. Towards this end, GO is tested through molecular simulations to understand the interactions between NDPK-B and NDPK-C. Influencing or modifying such enzyme active sites has been very less explored and, in this work, the molecular simulations suggest that GO is able to interact with the active site of NDPK-B to provide a sustained cAMP synthesis for longer duration. We found that conformational changes within NDPK-B and NDPK-C influence the interactions between them and such conformational changes are found to be governed by their adsorption on GO. Finally, we found the role of interfacial water molecules between NDPK-B and GO to be crucial in maintaining the interface between them.

## INTRODUCTION

In a recent report, American Heart Association announced that the prevalence of heart failure continues to rise over time, with aging of the population. An estimated 6.2 million American adults ≥20 years of age had heart failure between 2013 and 2016, compared with an estimated 5.7 million between 2009 and 2012^1^. Clearly, this is an ever-growing health concern, and research work regarding heart failure and especially molecular mechanisms underlying it are of increasing importance. Differences in molecular concentrations and activities account for the heart failure. Nucleoside diphosphate kinases (NDPKs) are a family of proteins involved in nucleotide homeostasis and protein histidine kinase^2^. There are four different types of NDPKs – A, B, C, and D. The ones involved with the G protein binding are the NDPK complex, NDPK-B and NDPK-C^3,4^. While similar, NDPK-B is a hexamer of identical chains and NDPK-C is a trimer of identical chains and the latter contains the amino acid tail that the NDPK-B does not have^5,6^. In a non-failing heart, the normal concentration of NDPK complex reacts with G_s_ (G stimulatory) protein, which causes second messenger cyclic adenosine monophosphate (cAMP) synthesis via the beta-adrenergic pathway. cAMP activates protein kinase A (PKA), which in turn activates the Ca^2+^ channels and maintains calcium homeostasis essential for normal cardiac function^7^. Failing heart, however, is associated with elevated NDPK and G_i_ (G inhibitory) contents. More NDPK binds with the G_i_ protein, which causes the suppression of cAMP synthesis and thus lower PKA activation, resulting in an imbalanced calcium ion homeostasis. This leads to heart failure when systolic and/or diastolic functions are limited^8,9^. G proteins (guanine-nucleotide binding proteins) are also a family of proteins that are membrane-bound signal transducers^10^. Two main types of G proteins – stimulatory and inhibitory – share a common heterotrimeric structure (containing □, β, and γ subunits). The □ subunit consists of the bound GDP molecule^11^. NDPK-B and in particular its residue Histidine-118 (His118) is involved in substrate channeling of phosphate to residue Histidine-266 (His266) on β subunit of G protein, which then releases G□ from the Gβγ complex, and the GDP on G□ accepts a phosphate from ATP thereby converting to GTP^3^. As a recent study suggests, NDPK-C is essential for the interaction between NDPK complex and G protein^4^. A possible therapeutic strategy against heart failure is to block the interactions between NDPK-BC and G_i_ protein. As the binding mechanism suggests, some options include blocking the His118 residue on NDPK-B or removing NDPK-C from the interaction scene. In this study, the investigated agent presumably capable of reaching these goals is graphene oxide (GO).

The choice of this nanomaterial was made because it is water-soluble and non-toxic^12–14^. It has also been shown in a recent article that GO has a potential to suppress the interactions between His118 and G protein by having NDPK-B adsorbed on GO^15^.

As NDPK-C proves to be essential for NDPK – G protein interaction whereas NDPK-B is capable of adsorbing on GO, in this article we investigate how GO affects the interactions of NDPK-B and NDPK-C. The existing scientific literature has little information to support the use of nanoparticles to suppress the interactions between NDPK-B and NDPK-C. Our hypothesis is that as NDPK-B adsorbs on GO, NDPK-C does not follow NDPK-B and is disassociated from it. Based on this it is proposed that GO is capable of removing NDPK-C from the interaction scene of NDPK complex and G protein. The objectives of this work are to quantify the molecular events between NDPK-B and NDPK-C in presence of GO and investigate how GO affects the interactions between them. These results are further compared with the control simulations between NDPK-B and NDPK-C in the absence of GO. Figure 1 shows the two systems with and without GO and the potential hypothesis along with the proposed effects based on the adsorption of NDPK-B on GO and its moving away from NDPK-C.

**Figure 1.**
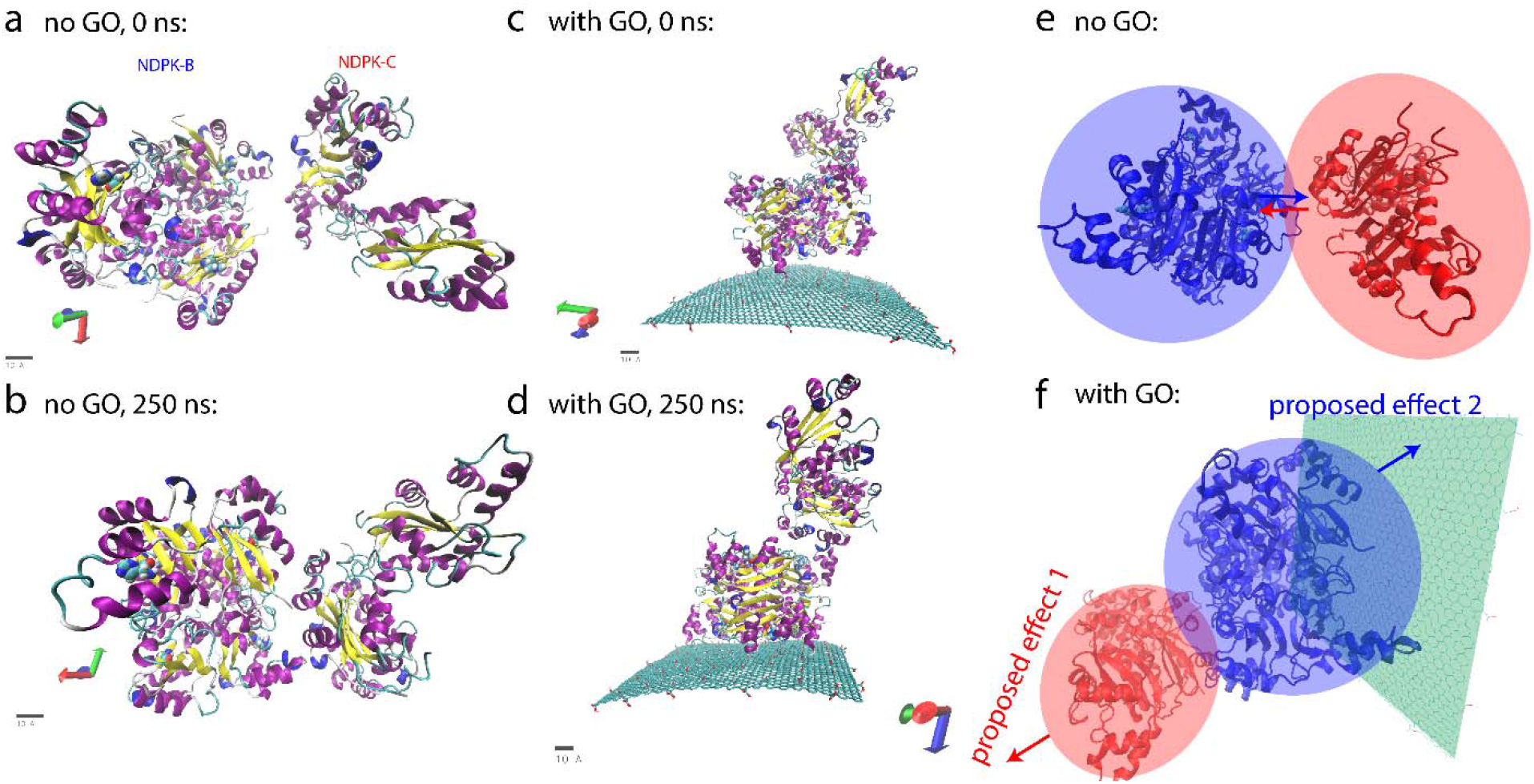
Trajectory screenshots and schemes of simulated systems: (a) — (d) Two systems (with and without GO) at the start (0 ns) and at the finish (250 ns) of the simulations. (e) — (f) Possible changes to the NDPK-BC complex that are hypothesized to occur based on the introduction of GO to the system. “Proposed effect 1” is the NDPK-C moving away from NDPK-B, “Proposed effect 2” is NDPK-B adsorbing on GO.

## RESULTS AND DISCUSSION

The trajectories obtained from the 250 ns production runs were analyzed based on the stability of the interacting molecules and energy evaluation of the interface. The first group is stability analysis, which includes root mean square deviation (RMSD), hydrogen bonds, center of mass (COM), salt bridges, and secondary structure analyses. Stability analysis typically determines how stable a molecule is within the timeframe of the simulation and its tendency to deviate, highlighting possible factors, both internal and external, influencing its behavior. The second group of analyses determined the various non-bonding energies of the molecules and is used to investigate the molecular adsorption tendencies evident from changes in energies. This group includes determining Van der Waals energies (between molecules) and electrostatic energies (both within and between molecules). Furthermore, analyses involving the optimal distance between NDPK-B and NDPK-C and between NDPK-B and GO were performed along with the role of interfacial water molecules during such interactions. We assign the time from the beginning of the simulation to ~140 ns as pre-interaction and from ~140 ns onwards as post-interaction.

As shown in Figures 2a and 2b, after 125ns, in the absence of GO, NDPK-B has a deviation of ~2 A□ whereas NDPK-C deviate by ~3.75 A□. In contrast to this, in the presence of GO, NDPK-B deviates by ~4 A□ while NDPK-C shows a deviation of ~3.6 A□ indicating that NDPK-B underwent significant conformational changes owing to its adsorption onto GO whereas NDPK-C started to lose its interaction with NDPK-B. From Figure 2c, the deviation of ~2.5 A□ before ~140 ns is indicative of the pre-interaction phase during which the interactions between NDPK-B and GO started to increase whereas from ~140ns onwards, there is an increase in the deviation of GO to ~3 A□, which indicates the post-interaction period during which not only the interactions between NDPK-B and GO became stronger but more importantly those between NDPK-B and NDPK-C would have started to diminish. These time checkpoints provide the preliminary time division of the pre- and post-interactions. The adsorption of NDPK-B onto GO and the eventual diminishing of the interactions between NDPK-B and NDPK-C is further verified by analyzing the energy of interaction between these molecules.

**Figure 2.**
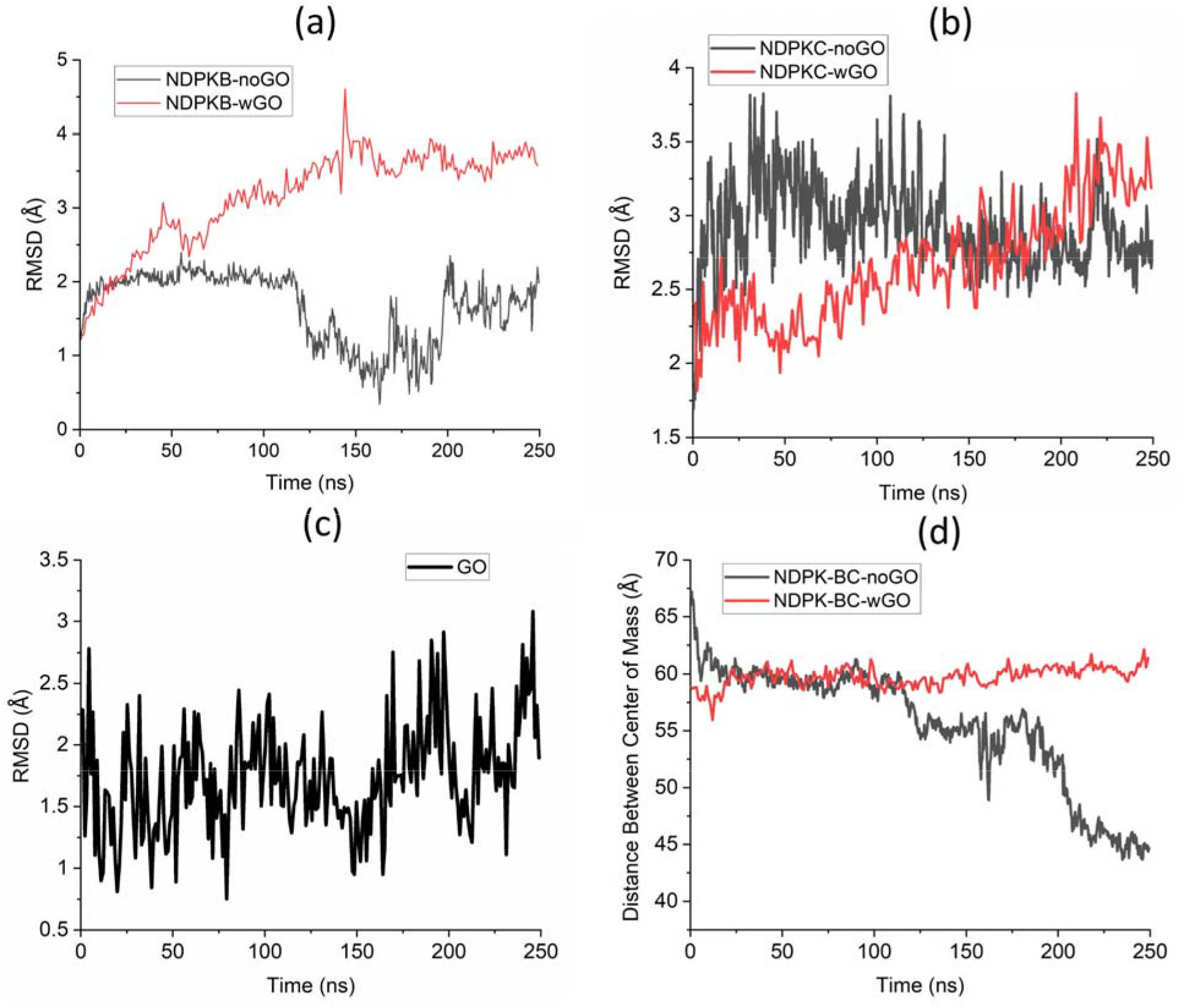
Stability Analysis of NDPK-B, NDPK-C in the presence and absence of graphene oxide. (a) RMSD of NDPK-B. (b) RMSD of NDPK-C. (c) RMSD of GO. (d) Distance between the center of masses (COM) of NDPK-B and NDPK-C. In all plots, noGO – no (absence of) GO; wGO – with (presence of) GO.

COM analysis utilizing the TCL scripts was used to investigate how distance between molecular centers of masses changes with respect to time, i.e., if different pairs of molecules move towards or away from each other. Figure 2d shows the COM distance between NDPK-B and NDPK-C in the presence and absence of GO and it is found that COM distance decreases starting at ~115 ns and further decreases to ~43 A□ beyond 200ns in the absence of GO, indicating that they come closer to each other as time goes by. On the contrary, when GO is present, initially, NDPK-B and NDPK-C do not change their positions relative to each other, whereas after ~20 ns, there is a jump in the COM data from 57 A□ to 60 A□. This is supported by the first milestone in the interactions in the presence of GO, where at ~50 ns, the interactions between NDPK-B and GO started to stabilize. Similarly, at ~150 ns and onwards, there is an increasing trend in the COM between NDPK-B and NDPK-C again supporting the second milestone when NDPK-B and NDPK-C started to lose interactions with each other. This finding indicates that the presence of GO refrains NDPK-B and NDPK-C from coming closer to each other. This may be interpreted as a possible way to block the interactions between these proteins.

Figure 3a shows that the distance between the center of masses of NDPK-B and GO decreases rapidly during the first 25 ns, clearly showing the two milestones at ~50 ns and ~150 ns, after which the distance between NDPK-B and GO reduces further, finally settling at ~30 A□, indicating an increase in the affinity interactions between NDPK-B and GO. Hydrogen bonds analysis was utilized to further confirm the stability of individual molecules and that of the interface between the molecules by quantifying the strength of interactions between them. A donor – acceptor distance of 3 A□ and the angle cut-off of 20 degrees is used as the criteria for the hydrogen bonds analysis. It is important to note that the hydrogen bond analysis do not include the salt bridges, which are analyzed separately. From Figure 3b, we can see that both in presence and absence of GO, the hydrogen bonds between NDPK-B and NDPK-C rapidly increase during the first 50 ns, then maintaining relatively stable values (though in the presence of GO, the average number is slightly larger). At around 200 ns, however, the number of hydrogen bonds starts to differ based on whether GO is present or not. In the presence of GO, the number of hydrogen bonds start to drop in comparison to the absence of GO, in which case, the hydrogen bonds rise substantially. This supports the proposed hypothesis that in the presence of GO NDPK-B and NDPK-C would experience diminished interactions. Non-bonding interaction energy analysis through Van der Waals (VDW) and electrostatic energies between NDPK-B and NDPK-C was done to understand the interactions at the interface. Figures 3c and 3d show the VDW interactions between NDPK-B and NDPK-C in the presence and absence of GO as well as those between NDPK-B and GO.

**Figure 3.**
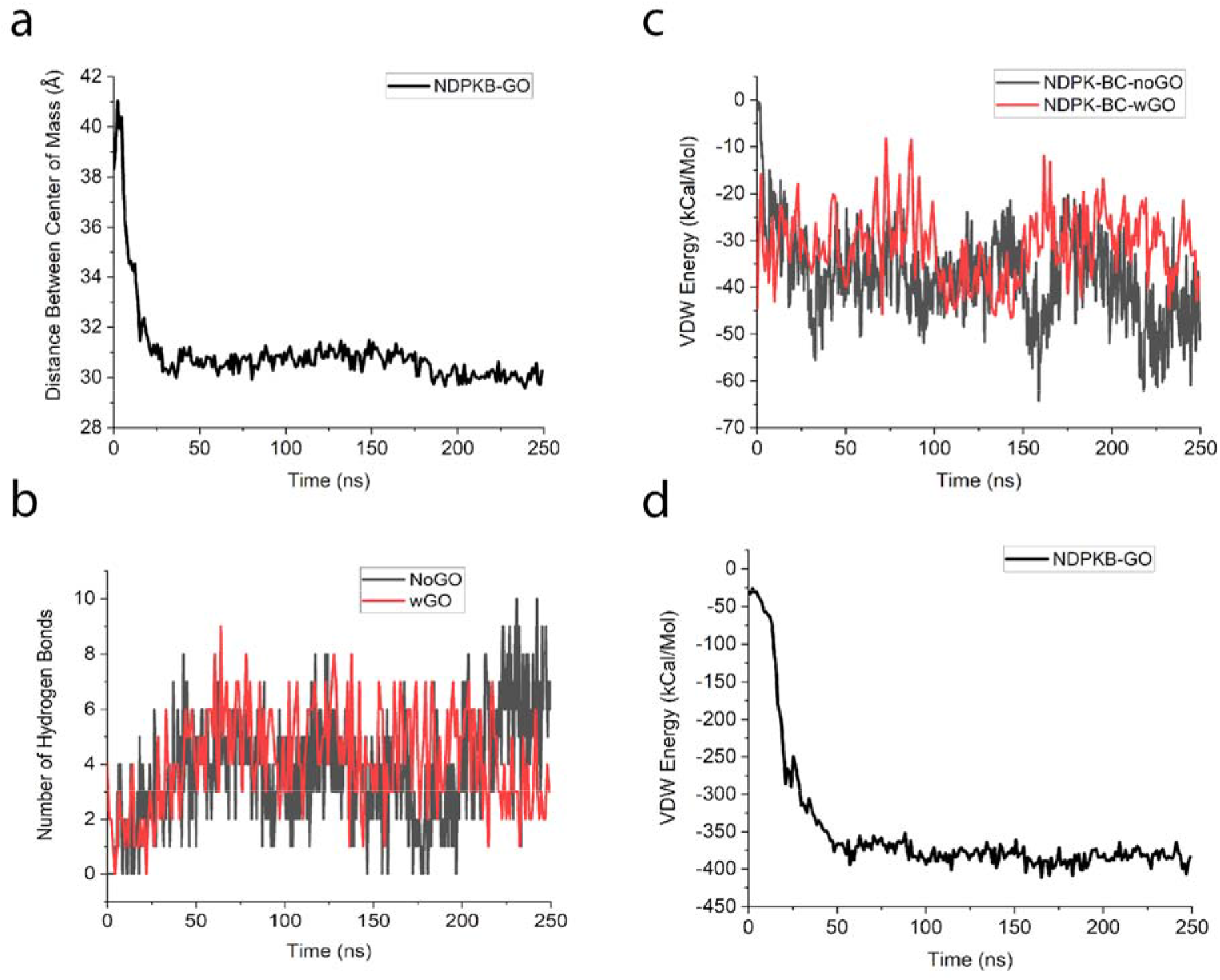
Stability and energetics analysis between NDPK-B, NDPK-C and graphene oxide. (a) Distance between the center of mass (COM) between NDPK-B and GO. (b) Number of hydrogen bonds between NDPK-B and NDPK-C in the presence and absence of GO. (c) Vander-Waals (VDW) energy between NDPK-B and NDPK-C in the presence and absence of GO. (d) VDW energy between NDPK-B and GO.

It is found that in the presence of GO, the VDW energy between NDPK-B and NDPK-C is comparable to that in the absence of GO, correctly pointing out that VDW energy is dominant largely in case of surface adsorption as evident from Figure 3d. As it can be seen from Figure 3d, the VDW energy between NDPK-B and GO is ~375 kCal/Mol at 50ns and after 150 ns, there is ~25 kcal/mol increase in the energy indicating that after 150 ns, NDPK-B started to lose its interactions with NDPK-C and got adsorbed on GO with larger energy.

As NDPK-B and NDPK-C interacts with each other, they would also form salt bridges to maintain the stable interactions. Table 1 shows the analysis of such intermolecular salt bridges between NDPK-B and NDPK-C. As we can see from the table, there are 12 salt bridges that are formed (< 4 Å distance between the interacting residues) between NDPK-B and NDPK-C throughout the course of the simulation in the absence of GO, while only two were formed in the presence of GO. In addition, four weak salt bridges (distance of 4 Å to 5 Å between the interacting residues) were detected without GO while only two such weak salt bridges were seen in the presence of GO. These two pieces of data combined imply that in the presence of GO, salt bridge formation between NDPK-B and NDPK-C is inhibited.

**Table 1.** Intermolecular salt bridges between NDPK-B and NDPK-C

| Salt bridge type | Without GO | With GO |
| --- | --- | --- |
| *Formed* | 12<br>ASP138—LYS49, ASP57—ARG152,<br>ASP57—ARG73, GLU141—LYS128,<br>GLU46—ARG105, GLU5—ARG73,<br>GLU67—LYS56, GLU71—LYS49,<br>GLU74—LYS66, GLU63—LYS85,<br>GLU62—LYS85, GLU152—ARG145 | 2<br>GLU152—ARG145, GLU62—LYS85 |
| *Weak* | 4<br>ASP121—LYS83, ASP57—ARG145,<br>GLU46—ARG102, ASP54—ARG152 | 2<br>ASP54—ARG152, GLU74—LYS124 |
| *Broken* | 3<br>GLU74—LYS124, GLU141—LYS49,<br>GLU141—LYS56 | 3<br>GLU5—ARG73, GLU63—LYS85,<br>ASP57—ARG152 |

Another aspect of interaction energy is based on electrostatic charges, which together with VDW energy quantifies the overall (total) interaction energy. Figure 4a shows electrostatic energy within NDPK-B, which is found to be more negative (attractive) in the presence of GO, starting 140 ns (post-interaction period). Similarly, for NDPK-C, in Figure 4b, it is found that the electrostatic energy is higher (more negative values) in the absence of GO, especially after 150ns. Interestingly, it is found that beyond 200ns, the overall electrostatic energy within NDPK-C in the presence of GO is very close to the absence of GO indicating that it regains its conformational structure once NDPK-B moves away. Together, these results indicate that the conformational energy landscapes for NDPK-B and NDPK-C are contrary to each other when it comes to adsorption. As can be seen, in the absence of GO, when NDPK-B interacts well with NDPK-C, it undergoes a sudden decrease (repulsion) in its conformational energy (Figure 4a) whereas NDPK-C undergoes a sudden increase (attraction) in its conformational energy (Figure 4b). This behavior related to the conformational energy is preserved in the presence of GO, where NDPK-B shows a decrease (repulsion) in its conformational energy but to a lesser extent due to its adsorption on GO.

**Figure 4.**
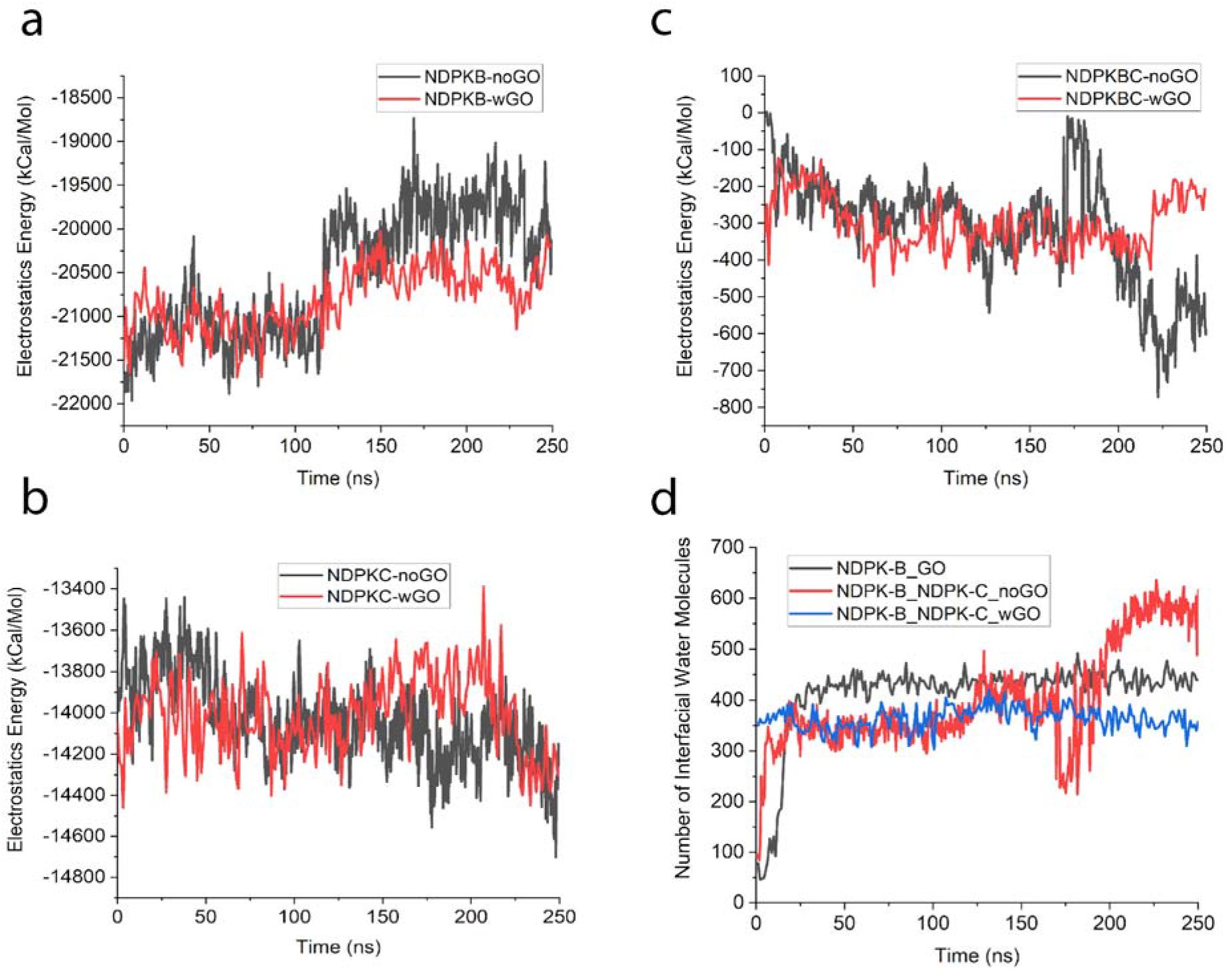
Electrostatics energy and role of interfacial water molecules. (a) Electrostatics energy within NDPK-B in the presence and absence of GO. (b) Electrostatics energy within NDPK-C in the presence and absence of GO. (c) Electrostatics energy between NDPK-B and NDPK-C in the presence and absence of GO. (d) Number of interfacial water molecules between NDPK-B and GO, and between NPDK-B and NDPK-C in the presence and absence of GO.

It is observed that this decrease in the electrostatic energy of NDPK-B in the presence of GO is because it interacts dominantly via the VDW energy with GO. Similar behavior is seen in NDPK-C where it shows a decrease in its conformational energy as it starts to lose its interactions with NDPK-B. Thus, it is found that in the event of adsorption, NDPK-B has a repulsive electrostatic effect whereas NDPK-C has an attractive electrostatic effect. Finally, comparing the electrostatic energy between NDPK-B and NDPK-C (Figure 4c), we find that in the absence of GO, both NDPK-B and NDPK-C are electrostatically bound by up to ~750 kcal/mol. Comparing this to the presence of GO, it is found that the maximum electrostatic energy between the two is at the most ~400 kcal/mol, which reduces further after 200 ns indicating that the interactions between NDPK-B and NDPK-C diminished significantly after 200ns.

To understand the interactions at the interface and the nature of the interface, both in the presence and absence of GO, a detailed analysis on the interfacial water molecules, their hydrogen bonds and RMSD was performed along with the optimal distance between the interacting molecules (NDPK-B and NDPK-C as well as NDPK-B and GO). As can be seen from Figure 4d, the number of interfacial water molecules between NDPK-B-GO complex as well as NDPK-B-NDPK-C complex are very different, where the number of interfacial water molecules (~450) in case of NDPK-B-GO complex is stable compared to those between NDPK-B-NDPK-C complex in the absence of GO, where the number of interfacial water molecules increases to ~600, especially after 200 ns (when interactions between NDPK-B and NDPK-C were stabilized). Figure 4d also shows that the number of water molecules at the interface of NDPK-B and GO increased sharply at 25 ns, when NDPK-B started to interact with GO, in the pre-interaction time and is then maintained at a relatively constant value. In contrast to this, the number of water molecules between NDPK-B and NDPK-C in the presence of GO were maintained at ~375 indicating the inhibiting role of GO to the interactions between NDPK-B and NDPK-C. This also further clarifies the role of interfacial water molecules in forming the complexes via hydrogen bonds. As can be seen from figures 5a and 5b, with a RMSD deviation of ~5 A□, the interfacial water molecules maintained on an average ~150 hydrogen bonds owing to the increased interactions between NDPK-B and GO starting at 25 ns.

**Figure 5.**
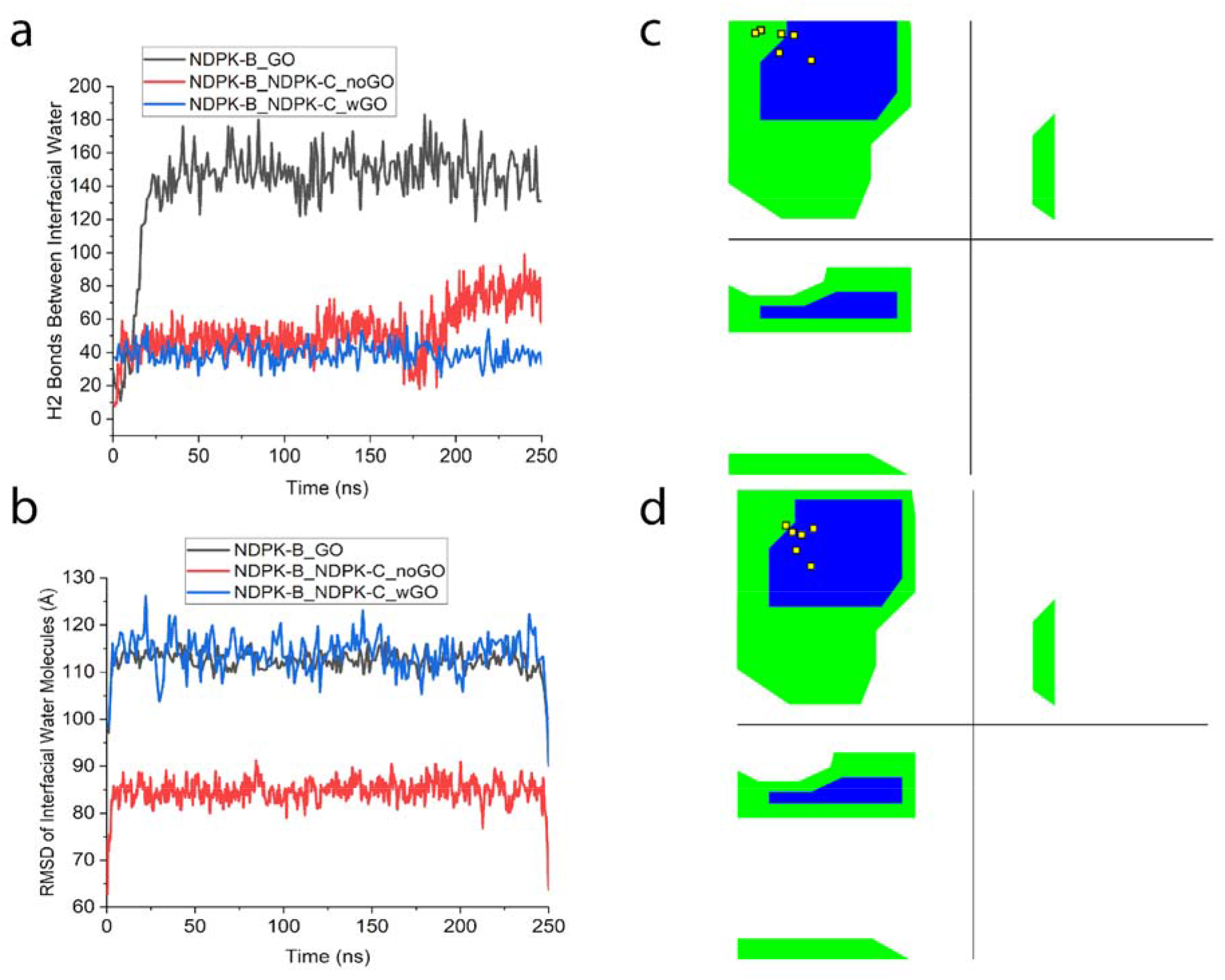
Regulation of interface between NDPK-B and NDPK-C by interfacial water molecules and Ramachandran plots. (a) Hydrogen bonds of the interfacial water molecules at the interface of NDPK-B and GO and between NPDK-B and NDPK-C in the presence and absence of GO. (b) RMSD of the interfacial water molecules between NDPK-B and GO, and between NDPK-B and NDPK-C in the presence and absence of GO. (c) – (d) Ramachandran plots for the six His118 residues in the absence and presence of GO.

A very different situation existed at the interface of NDPK-B and NDPK-C where comparing the presence and absence of GO, we find that in the absence of GO, the number of hydrogen bonds were found to be ~50 until 200 ns and ~80 after 200 ns when the interactions between NDPK-B and NDPK-C were stabilized with RMSD of ~8 A□, whereas in the presence of GO, the number of hydrogen bonds between NDPK-B and NDPK-C reduced to ~40 with an increased RMSD of ~15 A□. This analysis demonstrates that the presence of interfacial water molecules gives rise to better interactions at the interface with more water molecules present for stable interactions giving rise to more hydrogen bonds. Figures 5c and 5d present an additional layer of analyses of protein structures — the Ramachandran plots. They show the dihedral angles around alpha carbon atoms for the six His118 residues of NDPK-B in the absence (Figure 5c) and presence (Figure 5d) of GO. It is found that in the absence of GO, three of the six residues are in the blue (core) beta sheet region. Conversely, in the presence of GO, five of the six residues are in the blue region indicating that GO can influence the secondary structure around His118 residues. This finding is supported by the secondary structure analysis presented in the supplementary figure S4, where it is found that the residue His118 (part of the beta-sheet) undergoes significant conformational change in the presence of GO.

To determine the optimum distance between the interacting molecules, number of atoms and interaction energies were considered for the two complexes involving NDPK-B-NDPK-C as well as NPDK-B-GO. Furthermore, the effect of GO on NDPK-B-NDPK-C complex was also determined by comparing the optimum distance measurements in the presence and absence of GO. Figure 6 represents the first group of exhaustive analyses performed, aiming at finding the optimum distance between NDPK-B and GO. For both figures 6 and 7, a comparison was done with and without hydrogen atoms to understand the interactions between heavy atoms (oxygen, carbon, and nitrogen) as the heavy atoms are more stable and it was also useful to explore the effects of these heavy atoms on the interface.

**Figure 6.**
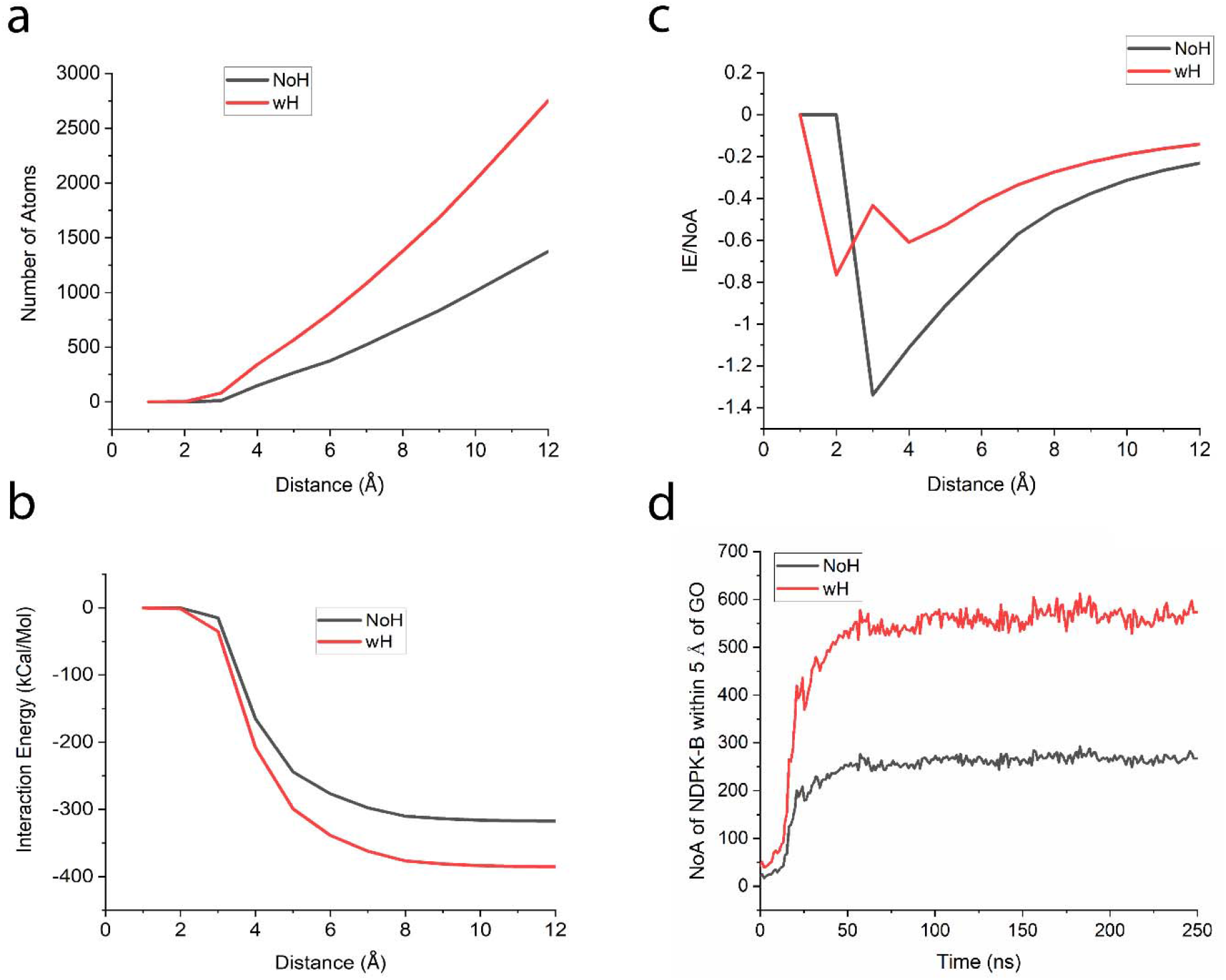
Optimal distance analysis between NDPK-B and GO. (a) Number of atoms between NDPK-B and GO as a function of distance with and without hydrogen atoms. (b) Total interaction energy between NDPK-B and GO as a function of distance with and without hydrogen atoms. (c) Maximum interaction energy per atom between NDPK-B and GO as a function of distance with and without hydrogen atoms. (d) Number of atoms (NoA) of NDPK-B within 5 Å of GO as a function of time with and without hydrogen atoms.

**Figure 7.**
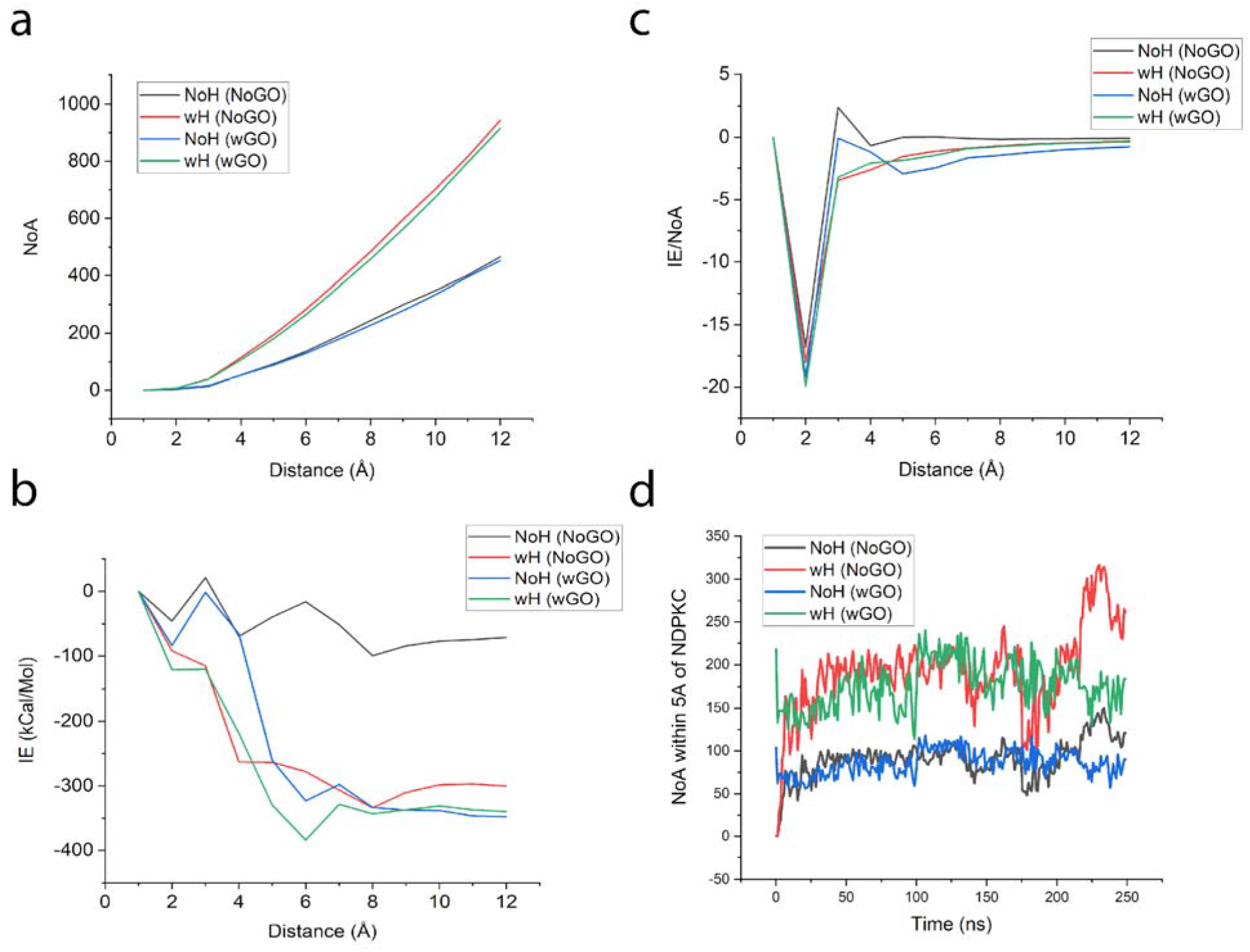
Optimal distance analysis between NDPK-B and NDPK-C. (a) Number of atoms between NDPK-B and NDPK-C as a function of distance in the presence and absence of GO with and without hydrogen atoms. (b) Interaction energy between NDPK-B and NDPK-C as a function of distance in the presence and absence of GO with and without hydrogen atoms. (c) Maximum interaction energy per atom between NDPK-B and NDPK-C in the presence and absence of GO with and without hydrogen atoms. (d) Number of atoms (NoA) of NDPK-B within 5 Å of NDPK-C with respect to time in the presence and absence of GO with and without hydrogen atoms.

All the values were collected from the time the system became stabilized at ~50 ns. This timestamp was obtained by looking at the stability from the RMSD plot. Figure 6a shows how the number of atoms of NDPK-B (average number with and without hydrogen atoms) is increasing as the distance from the surface of GO was increased from 1 A□ to 12 A□ (non-bonding cut-off limit). This is not surprising as we continue to increase the distance from GO, we anticipate encountering more NDPK-B atoms owing to an increase in the radius of our spherical “search spotlight”. Similarly, in Figure 6b, the values of the interaction energy, IE (non-bonding energies as a sum of VDW and electrostatic energies), of NDPK-B atoms are decreasing as the spherical “search spotlight” radius centered at GO is increasing from 1 A□ to 12 A□. As more NDPK-B atoms are included in the spherical selection zone, we find that beyond ~8 A□, the interactions energy flattens out indicating that there are ~600 heavy atoms (without hydrogen) that play a crucial role in maintaining the interface between NPDK-B and GO. Finally, to determine the optimal distance between NDPK-B and GO, interaction energy per atom is determined as a ratio of IE/NoA for the respective distances (Figure 6c). The negative peak indicates the optimal adsorption distance between NDPK-B and GO of ~3 A□. This further also indicates that ~600 atoms of NDPK-B adsorbed on the surface of GO are enough to maintain the stability of the interface. Figure 6d illustrates a more in-depth look at the number of atoms of NDPK-B within 5 A□ of GO throughout the whole course of the simulation. It shows that the number of atoms increased over a period, and after stabilization was achieved, maintained a relatively constant value of ~275 (without hydrogen) and ~575 (with hydrogen) atoms. This also adds an additional lens to show that NDPK-B and GO interacted favorably starting ~50ns and maintained a consistent number of atoms within the adsorption limits.

On similar lines, Figure 7 represents exhaustive analysis performed on the interface between NDPK-B and NDPK-C to understand and compare the nature of this interface to that of NDPK-B-GO. Figure 7a shows a comparison between the number of atoms of NDPK-B at certain distances from NDPK-C in the presence and absence of GO. It is found that the number of atoms in the presence and absence of GO, though slightly more in the absence of GO, are very similar.

For the determination of optimal distance between NDPK-B and NDPK-C, as can be seen from Figures 7b and 7c, it is 2 A□ regardless of the presence or absence of GO, which is less than the optimal distance between NDPK-B and GO pointing out the differences in the interactions between the adsorption of NDPK-B on GO, which is largely due to the non-binding energies such as Van der Waals, whereas that between NDPK-B and NDPK-C is largely due to electrostatic interactions. Finally, over a period of 250 ns, the number of atoms of NDPK-B within 5 A□ of NDPK-C in the presence and absence of GO shows the dynamics of the interface between NDPK-B and NDPK-C as shown in Figure 7d providing important insights into the early onset of binding between NDPK-B and NDPK-C, after 200 ns, in the absence of GO as can be seen from the comparison between heavy atoms (NoH, NoGO) and (NoH, wGO) plots.

## MATERIALS AND METHODS

The two systems (NDPK-BC alone and NDPK-BC with GO) were first modeled in Visual Molecular Dynamics (VMD)^16^, a software used to display and analyze biomolecular systems. The PDB files for NDPK-B and NDPK-C were obtained from Protein Data Bank^17^ (codes 1NUE and 1ZS6, respectively)^5^, which were then used to generate the protein structure files (PSF) using the CHARMM (Chemistry at HARvard Macromolecular Mechanics) force field parameter files^18^. GO was created using the Molefacture plugin in VMD from the graphene sheet generated using an inbuilt graphene sheet builder plugin, which was then used to create the GO flake of size 96 A□ × 90 A□. The chemical structure of the flakes used is C10O1(OH)1(COOH)0.5, also known as OGO^19^. To merge GO with NDPK-BC to set up the second combined system, TCL scripts for moving and rotation of GO were employed. GO was intentionally placed on one side of NDPK-B (~10 Å from its active sites, His118), with NDPK-C on the other side of NDPK-B to minimize the concomitant interactions between NDPK-C and GO. The systems were solvated using the Transferable Intermolecular Potential with 3 Points (TIP3P) water model^20^ using the built-in Solvate plugin in VMD. The resulting water box was set up so that each box edge was 10 □ away from the nearest atom. Solvated system was neutralized with NaCl to compensate for the overall net charge in the system, resulting in a total of ~134,000 atoms for the NDPK-BC system and ~382,000 atoms for the NDPK-BC+GO combined system. All simulations were carried out using Nanoscale Molecular Dynamics (NAMD)^21^ on an Intel Core i9 cluster with a total of 36 cores, and NVIDIA GeForce RTX 2080 GPU. The temperature was maintained at 310 K by the Langevin Thermostat and the pressure was set to 1 atm using Nose-Hoover Langevin-piston barostat with a period of 100 ps and a decay rate of 50 ps. A cut-off of 10–12 A□ was used for short-range forces and particle mesh Ewald algorithm was used for calculating long-range forces with a time step of 2 fs. A 5000-step energy minimization was performed first to reach a minimum for the potential energy followed by an equilibration of 500,000 steps (1 ns). Production runs were performed for 250 ns for each simulation. Analysis of the generated molecular trajectories was primarily conducted in VMD, with the help of TCL and MATLAB scripts. Trajectory analysis was performed using VMD plugins for hydrogen bonds, RMSD, non-bonding and conformational energies, Ramachandran Plot, and salt bridges, as well as the external TCL scripts to analyze distance between center of mass (COM) and number of atoms (NoA). Salt bridges data was further analyzed using a MATLAB script, with cutoff distance of 4 □.

## CONCLUSION

In this work, the behavior of NDPK-B/C complex was assessed in the absence and presence of GO. It was found that GO causes NDPK-B to adsorb, change conformation, and thus lose interactions with NDPK-C. This was observed through a reduced COM distance between NDPK-B and GO (from 40 A□ to 30 A□) and increasing COM distance (from 50 A□ to 60 A□) between NDPK-B and NDPK-C. Furthermore, with an optimal distance of 2 A□, the interactions between NDPK-B and NDPK-C were found to be largely governed by electrostatic energies (300 kcal/mol) whereas those between NDPK-B and GO were found to be largely governed by Van der Waals energies (375 kcal/mol) as evident from the optimal distance of 3 A□. It was also found that the number of water molecules at the interface of NDPK-B and GO (~425 molecules) as well as NDPK-B and NDPK-C (~600 molecules in the absence of GO and ~375 molecules in the presence of GO) played a crucial role for the adsorption and that the water molecules formed increased hydrogen bonds only in the presence of favorable interactions at the interface. Diminished interactions within the NDPK-B/C complex can reduce its ability to phosphorylate GDP(i), therefore GO can have an impact of diminishing NDPK-B/C activity in the pathways inhibiting the cAMP formation.

## Supporting information

Supporting Information: Data and Software Sharing

Author Contributions

## ASSOCIATED CONTENT

### Supplementary Information

Screenshots of the 250ns trajectories of the NDPK-B and NDPK-C interactions in the presence and absence of GO in figures S1 and S2, number of intramolecular hydrogen bonds within NDPK-B and NDPK-C in figure S3, Secondary structure analysis of His118 and its neighboring residues in the presence and absence of GO in figure S4, and Tables ST1 and ST2 showing the intramolecular salt bridges within NDPK-B and NDPK-C.

### Data and Software Availability

Modeling and Simulation software, VMD (Visual Molecular Dynamics) and NAMD (Nanoscale Molecular Dynamics) used in this study are freely available from the Theoretical and Computational Biophysics group at the NIH Center for Macromolecular Modeling and Bioinformatics at the University of Illinois at Urbana-Champaign, http://www.ks.uiuc.edu/Research/vmd/

Atomic coordinate (protein data bank, PDB) files for the two enzymes, NDPK-B and NDPK-C, 1NUE and 1ZS6 were obtained from the database located at Research Collaboratory for Structural Bioinformatics (RCSB), www.rcsb.org.

To create the protein structure files (PSF) and to carry out all – atom simulations, the necessary topology and forcefield parameter files were obtained from the Chemistry at Harvard Macromolecular Mechanics (CHARMM) database located at the MacKerell Lab at the University of Maryland, School of Pharmacy, http://mackerell.umaryland.edu/charmm_ff.shtml. The models for the two systems with and without graphene oxide and the two enzymes NDPK-B and NDPK-C along with the parameter and configuration files are also available as part of the supplementary information.

The supplementary data contains a README text file describing the different data files related to modeling and simulation of the systems, as well as the data files themselves.

## AUTHOR INFORMATION

The authors declare no competing financial interest.

## ACKNOWLEDGMENT

We acknowledge Fairfield University for providing computational resources.

## ABBREVIATIONS

NDPK: nucleoside diphosphate kinase
GO: graphene oxide
VMD: Visual Molecular Dynamics
NAMD: NAnoscale Molecular Dynamics

