## Supplementary Figures for "Role of graphene oxide in inhibiting the interactions between nucleoside diphosphate kinases -B and -C"

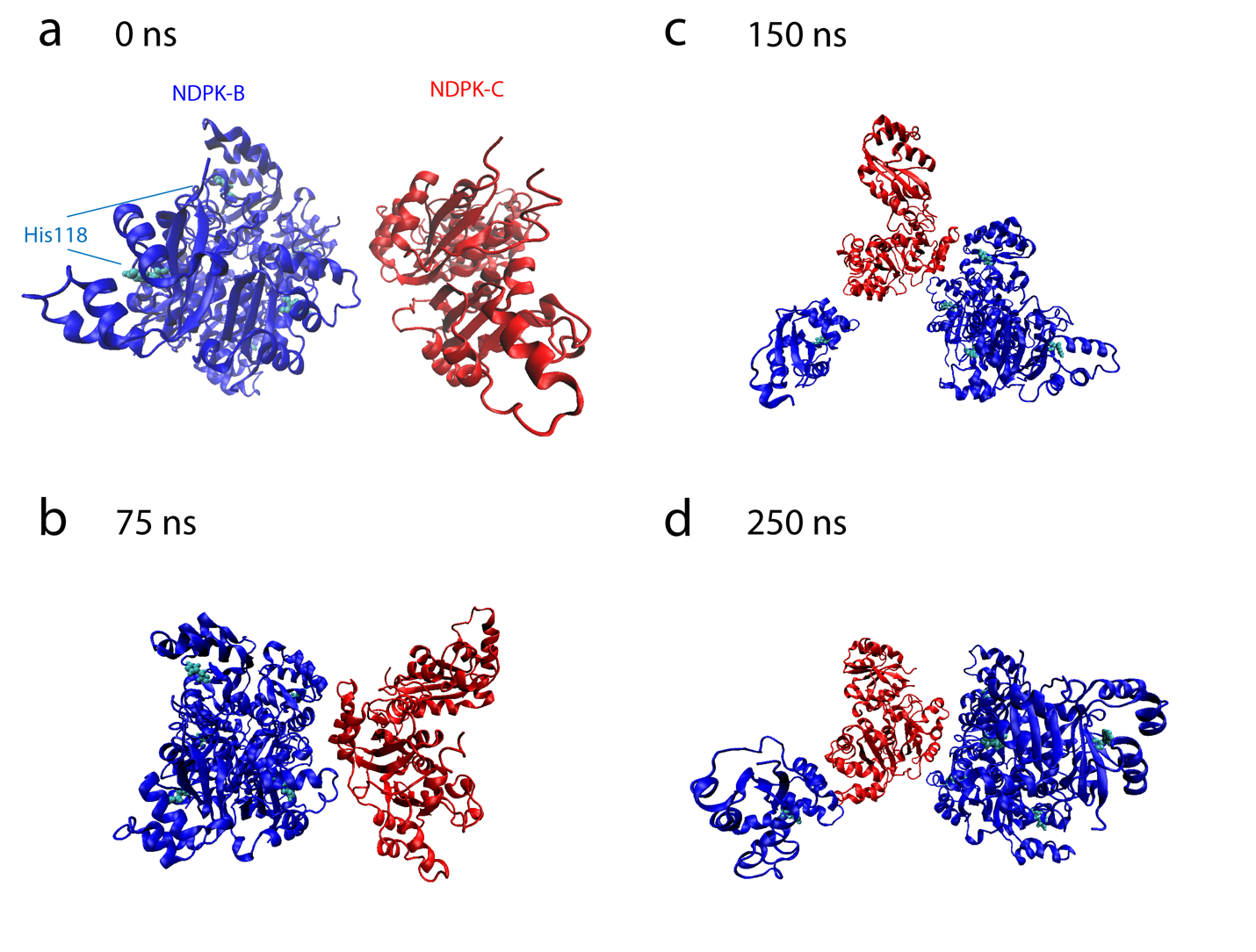

**Figure S1.** **Screenshots of the NDPK-BC system at four points throughout the simulation:** (a) 0 ns; (b) 75 ns; (c) 150 ns; (d) 250 ns. The position of the observer remains the same for all four screenshots.

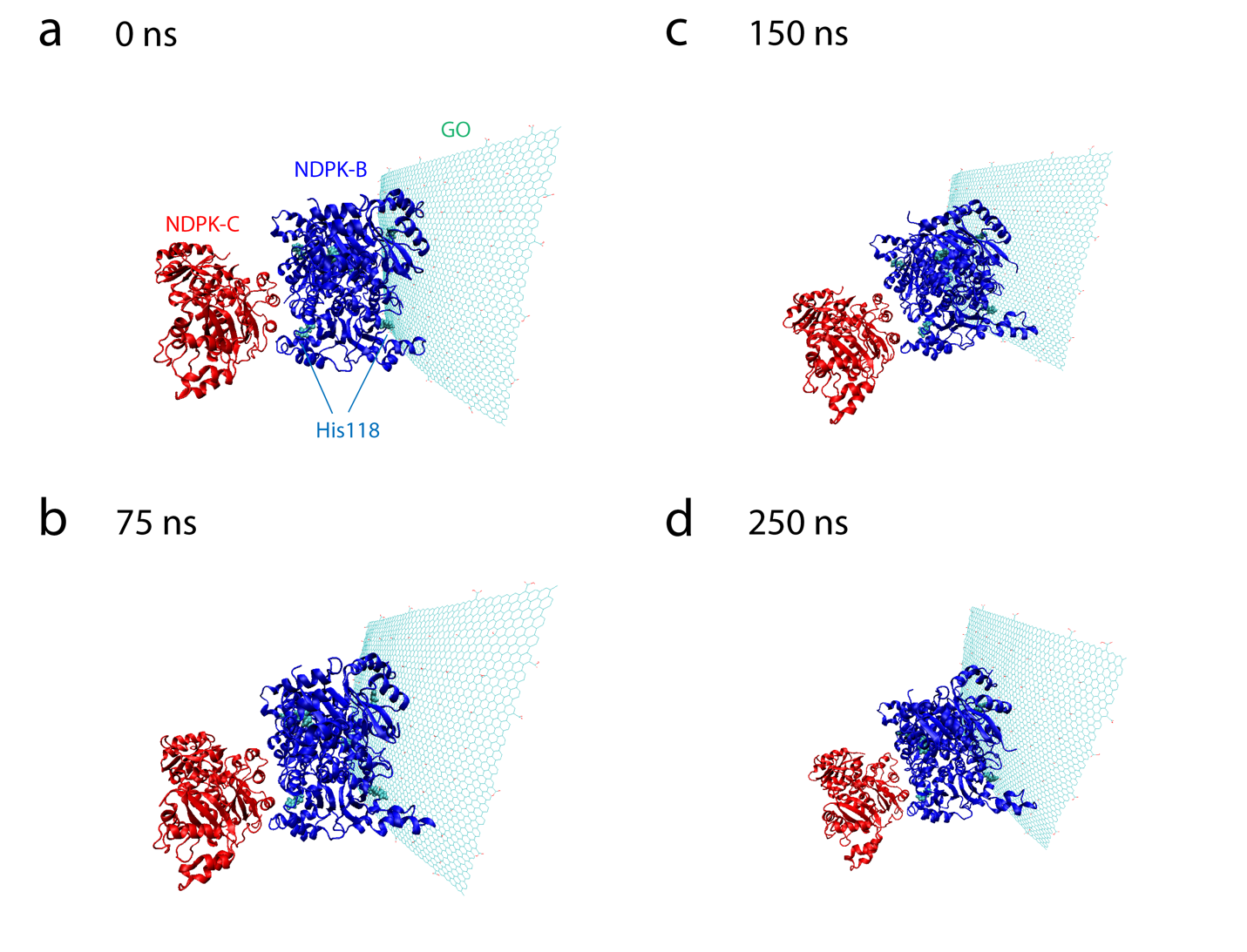

**Figure S2.** **Screenshots of the NDPK-BC system in the presence of GO at four points throughout the simulation:** (a) 0 ns; (b) 75 ns; (c) 150 ns; (d) 250 ns. The position of the observer remains the same for all four screenshots.

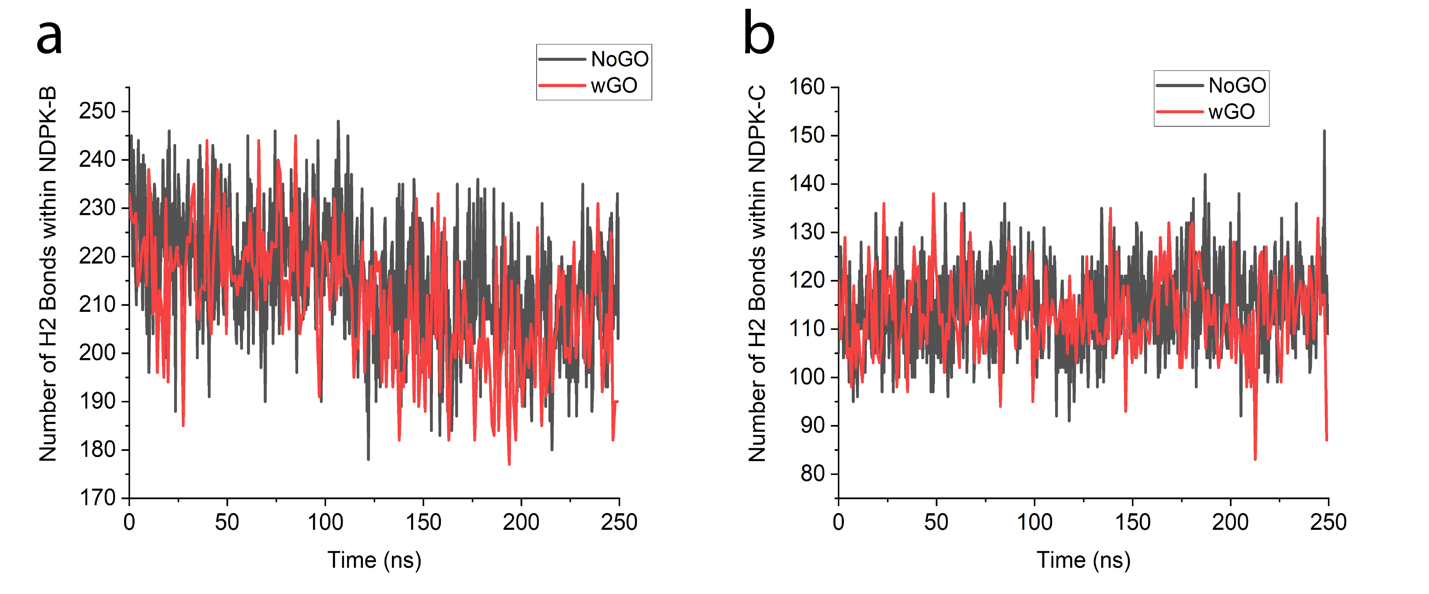

**Figure S3.** **Number of intramolecular hydrogen bonds within:** (a) NDPK-B; (b) NDPK-C.

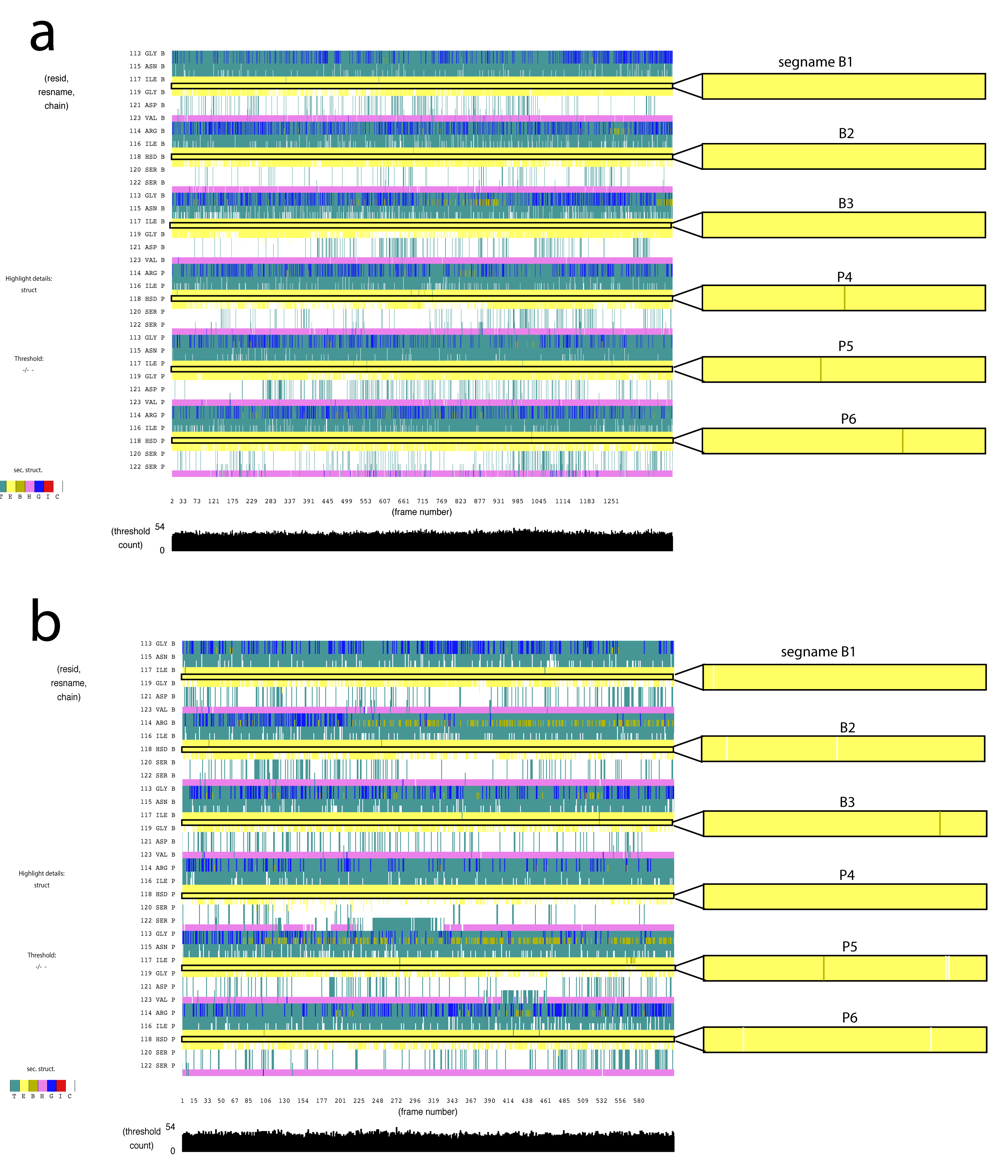

**Figure S4. Secondary structure analysis of six His118 residues of NDPK-B measured along with 5 residues in vicinity, zoomed in on His118:** (a) in the absence of GO; (b) in the presence of GO.
