## Supporting Information: Data and Software Sharing for "Role of graphene oxide in inhibiting the interactions between nucleoside diphosphate kinases -B and -C": README.txt

The supplementary data zip file contains the following data files:

For NDPK-BC control simulations:
- ionized.pdb: PDB file of NDPK-B, NDPK-C, water, and ions
- ionized.psf: PSF file of NDPK-B, NDPK-C, water, and ions
- ndpkBC.conf: configuration file for carrying out NAMD simulations
- par_all22_prot.prm: CHARMM parameter file for proteins

For NDPK-BC-GO simulations:
- NDPK-BC+GO_ionized.pdb: PDB file of NDPK-B, NDPK-C, GO, water, and ions
- NDPK-BC+GO_ionized.psf: PSF file of NDPK-B, NDPK-C, GO, water, and ions
- GO.prm: parameter file for GO
- Prod.conf: configuration file for carrying out NAMD simulations
- par_all22_prot.prm: CHARMM parameter file for proteins
- toppar_water_ions.str: parameter file for water and ions
