## Supplementary material for "Role of graphene oxide in inhibiting the interactions between nucleoside diphosphate kinases -B and -C": Author Contributions

IM conceptualized and designed the study. AZ modeled the control and experimental systems using VMD. IM carried out the NAMD simulations. AZ performed the formal analysis on the simulated trajectories. IM did the data curation and funding acquisition for the project. IM did the project administration and provided the resources. IM did the supervision of the project and validation of the acquired results. AZ prepared the original draft for the manuscript. IM reviewed and edited the manuscript. All authors approved the submitted version of the manuscript and agree to the author list and author contributions.
